# Genetic transformation and live-cell nuclear and actin dynamics during the life cycle of a chytrid

**DOI:** 10.1101/787945

**Authors:** Edgar M. Medina, Kristyn A. Robinson, Kimberly Bellingham-Johnstun, Giuseppe Ianiri, Caroline Laplante, Lillian K. Fritz-Laylin, Nicolas E. Buchler

## Abstract

Chytrids are early-diverging fungi that share ancestral features of animals, including cells that crawl and swim. At later stages, chytrid cells resemble fungi with a chitin-based cell wall and hyphal-like structures known as rhizoids. Chytrids are important evolutionary transitional forms, but much remains unknown about their cell biology because we lack genetic tools for the live-cell imaging of their nuclear and cytoskeletal dynamics. Here, we generated stable transgenic lines of the soil chytrid *Spizellomyces punctatus*, and coupled live-cell microscopy and fluorescent tagging to measure the timing and coordination of growth, the cell cycle, and the actin cytoskeleton. We show that *Spizellomyces* zoospores rapidly encyst, develop rhizoids, and undergo multiple rounds of synchronous nuclear division in a sporangium, followed by cellularization, to create and release hundreds of zoospores. The life cycle is complete in less than 30 hours. We further demonstrate that crawling zoospores, akin to animal cells, display polymerized actin at the leading edge of amoeboid fronts. After encystment, polymerized actin reorganizes into fungal-like cortical patches and cables that extend into the rhizoid. Actin remains highly dynamic during sporogenesis with the formation of actin perinuclear shells each cell cycle and the emergence of polygonal territories during cellularization. *Spizellomyces* is a fast-growing and genetically-tractable organism that should be useful for comparative cell biology and understanding the evolution of fungi and early eukaryotes.

## Introduction

Zoosporic fungi, commonly referred to as “chytrids”, span some of the deepest fungal Phyla and comprise much of the undescribed environmental fungal DNA diversity in aquatic ecosystems (1–4). Many chytrids are saprophytes or parasites of photosynthetic organisms and actively shuttle carbon to higher trophic levels (4, 5). Other chytrids are animal parasites, including the infamous *Batrachochytrium* genus that includes the frog-killing B. *dendrobatidis* (6) and salamander-killing B. *salamandrivorans* that are devastating global amphibian populations (7).

Chytrids are unique in that they have retained ancestral cellular features, shared by animal cells and amoebae, while also having fungal features. For example, chytrids begin their life as motile zoospores that lack a cell wall, swim with a single posterior cilium nucleated from a centriole, and crawl across surfaces (8–12). Later life cycle stages exhibit fungal characteristics including the formation of chitinous cell walls, the growth of hyphal-like structures, and the development of a sporangium (sporangiogenesis); see Fig. 1. Chytrid zoosporogenesis involves multiple rounds of mitosis without cytokinesis to create a multi-nuclear coenocyte, followed by cellularization to form zoospores with a single nucleus (Fig. 1). The formation of a multinuclear compartment followed by cellularization is reminiscent of development in flies (e.g. *Drosophila*), amoeba (e.g. *Physarum*), and protozoa (e.g. *Plasmodium*). Although there are important differences (particularly the need for the chytrid sporangium to extract nutrients from the environment and coordinate growth with the cell cycle), determining the mechanisms controlling chytrid cellularization provides a comparative framework for understanding cellularization in animals and other eukaryotic lineages.

**Fig. 1.**
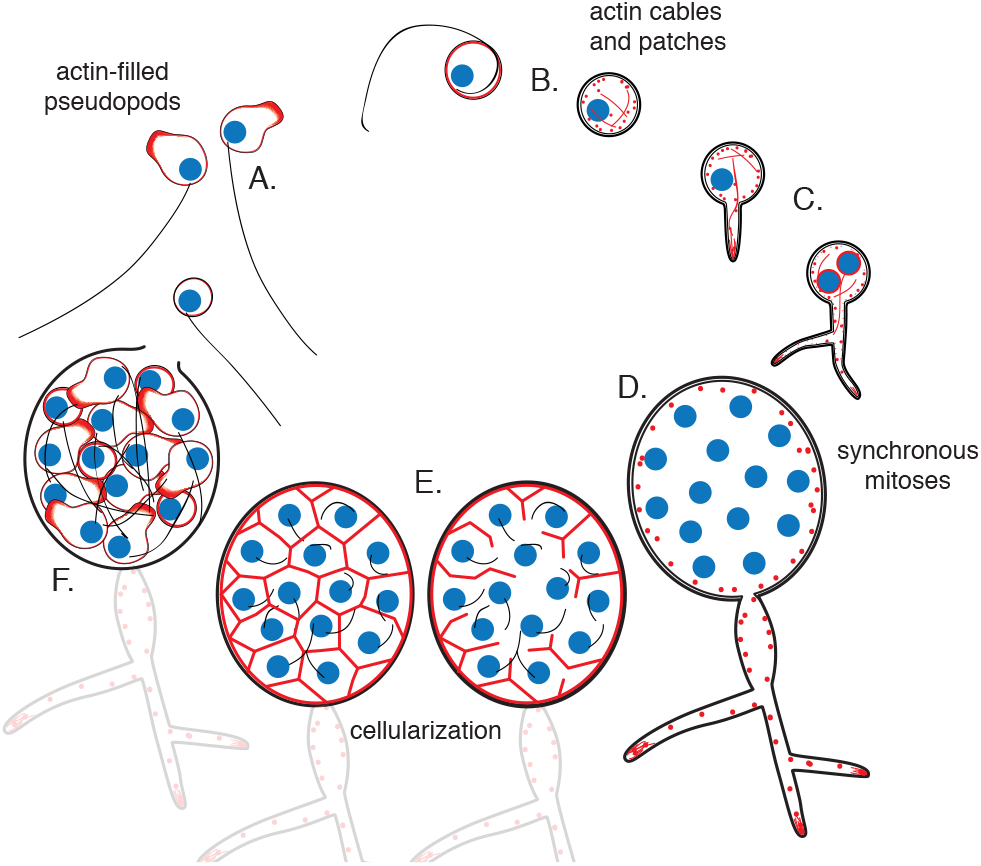
Life cycle of the chytrid *Spizellomyces punctatus*. Timeline and events were measured in this work. This chytrid produces globular zoospores (3-5 μm) that swim with a whip-like motile cilium (20-24 μm). **A)** The uninucleated zoospore (nucleus in blue) has a cilium associated with a basal body. Swimming zoospores can also crawl on surfaces using amoeboid-like motion (polymerized actin in red). **B)** At the start of encystment, the cilium usually retracts by a lash-around mechanism followed by formation of cell wall (14). **C)** The cyst then germinates and forms a single germ tube that later expands and branches into a rhizoidal system. The nucleus remains in the cyst during germ tube expansion as the cyst develops into a single reproductive structure called the sporangium. The first mitotic event correlates with ramification of rhizoids from the germ tube. **D)** Mitosis in the sporangium is coordinated with growth, as nuclei replicate and divide in a shared compartment. There can be a total of 4 to 8 synchronous mitotic cycles as each sporangium develops a branched rhizoid system with subsporangial swelling in the main rhizoid. **E)** Mitosis halts and zoospore formation begins in the sporangium. Ciliogenesis likely occurs before cellularization (*Allomyces*, (15)) **F)** The nuclei cellularize and develop into zoospores while the sporangium develops discharge papillae. Once cellularization is complete and environmental conditions are favorable, the zoospores will escape the sporangium through the discharge papillae. Diagram not drawn to scale.

The major bottleneck to studying chytrids in molecular detail has been the lack of a model organism with tools for genetic transformation. Here, we describe the successful adaptation of *Agrobacterium*-mediated transformation to generate reliable and stable genetic transformation of the soil chytrid *Spizellomyces punctatus*. We expressed fluorescent proteins fused to histone and actin-binding proteins to characterize the development and cell biology of *Spizellomyces* throughout its life cycle using live-cell imaging. Below, we show that *Spizellomyces* is a well-suited model system for uncovering molecular mechanisms of cell cycle regulation, cell motility, and development because it is fast-growing, displays both crawling and swimming motility, and possesses a characteristic chytrid developmental life cycle. These tools allow, for the first time, direct molecular probing to test hypotheses about the evolution and regulation of the cell cycle (13), cell motility (12), and development in chytrid fungi.

## Results

### Developing tools for genetic transformation

The plant pathogen *Agrobacterium tumefasciens* normally induces plant tumors by injecting and integrating a segment of transfer DNA (T-DNA) from a tumor-inducing plasmid (Ti-plasmid) into the plant genome. Researchers have exploited this feature to integrate foreign genes in plants by cloning them into the T-DNA region of the Ti-plasmid, inducing virulence genes for processing/transport of T-DNA, and co-culturing induced *Agrobacterium* with the desired plant strain. Because Agrobacterium-mediated transformation has been adapted for transformation of diverse animals and fungi (16–20), we chose to use this system in *Spizellomyces punctatus*. To this end we modified an *Agrobacterium* plasmid to integrate and express a selectable marker (e.g. drug resistance) in *Spizellomyces*. To determine a suitable selection marker for *Spizellomyces*, we tested the effects of drugs on the growth of the chytrid on agar plates. We spread zoospores on plates with various concentrations of drugs and assessed the cultures for cell growth, colony formation, and zoospore release using light microscopy. Although Geneticin (G418), Puromycin, and Phleomycin D10 (Zeocin) did not inhibit growth up to 800mg/L, we determined that 200mg/L Hygromycin and 800mg/L Nourseothricin (CloNAT) resulted in complete absence of growth after 6 days of incubation at 30°C. All remaining experiments were performed using Hygromycin (200mg/L).

Next, we identified *Spizellomyces* promoters that can drive gene expression at enough levels to provide resistance to hygromycin and measurable protein fluorescence. In the absence of a chytrid system to perform these tests, we reasoned that *Spizellomyces* promoters that express at high levels in yeast (*Saccharomyces cerevisiae*) would likely also work in chytrids. Therefore, we used an *Agrobacterium* plasmid (19) that propagates in yeast to first screen *Spizellomyces* promoters that successfully express a fusion of hygromycin resistance (hph) and green fluorescent protein (GFP); see Materials & Methods. We confirmed that *Spizellomyces* HSP70 and H2B promoters resulted in resistance to hygromycin as well as measurable GFP fluorescence in yeast via flow cytometry (Fig S1A). All remaining experiments were performed using the stronger H2B promoter.

With active promoters and effective selection parameters in hand, we performed *Agrobacterium*-mediated transformation by co-culturing *Spizellomyces* zoospores with *Agrobacterium* carrying H2Bpr-hph-GFP; see Materials & Methods methods. Although hygromycin-resistant, none of the transformants exhibited green fluorescence above background (Fig. S2A). When GFP was replaced by tdTomato, we obtained transformants that exhibited both hygromycin resistance (Fig. S2A) and cytoplasmic fluorescence (Fig S2B). Further tests with other fluorescent proteins showed that mClover3, mCitrine, and mCerulean3 are functional in *Spizellomyces* (Fig. S3). Once we confirmed that both selection marker and fluorescent protein were functional in *Spizellomyces*, we designed a construct with greater applicability, in which these markers are functionally independent and the fluorescent protein (tdTomato) is fused in-frame to the C-terminal of a protein of interest (POI). This design exploits the compact and divergent architecture of *Spizellomyces* H2A/H2B promoters to express (POI)-tdTomato in an upstream direction (H2B promoter) while expressing hph in an downstream direction (H2A promoter). As a proof of concept and because we were interested in following nuclear dynamics to measure the timing and synchrony of mitotic events (e.g. DNA segregation, see next section), our first protein of interest was histone H2B; see Fig. 2A.

**Fig. 2.**
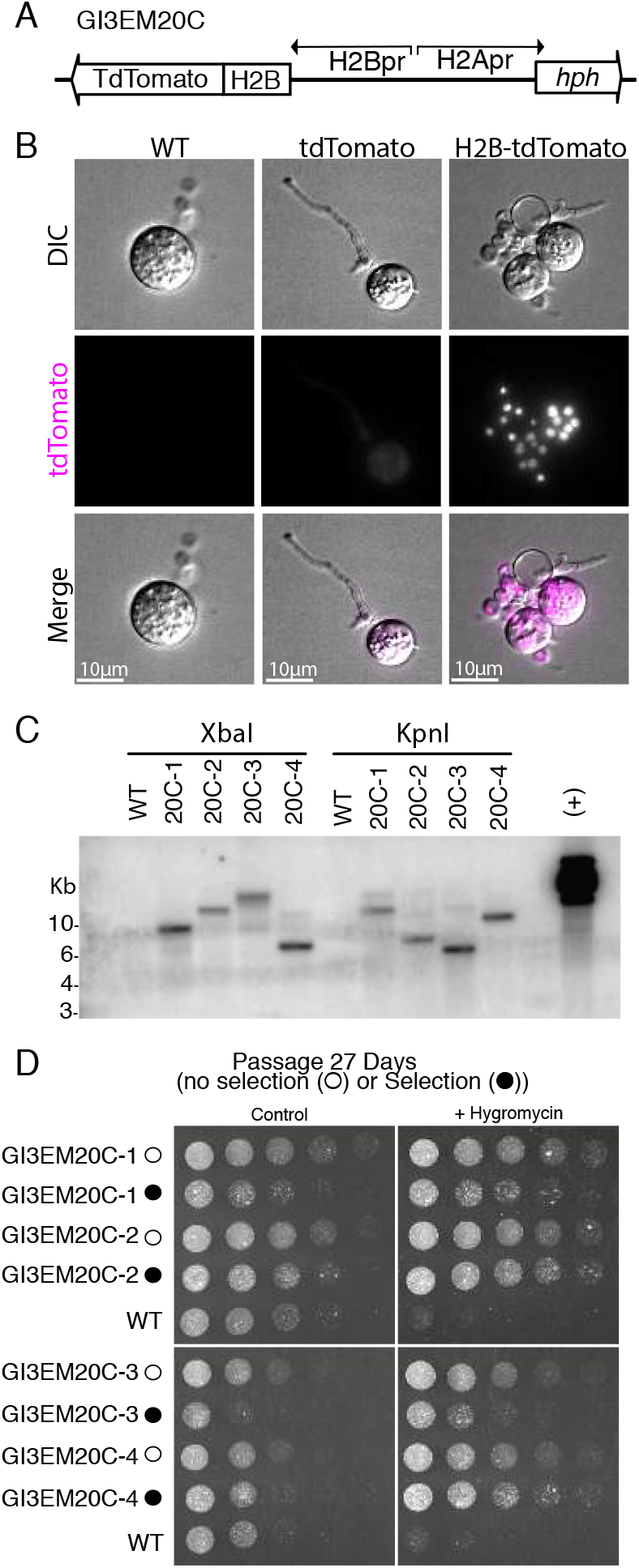
Genomic integration of H2B-tdTomato using *Agrobacterium-mediated* transformation. **A**) Plasmid GI3EM20C takes advantage of the divergent architecture of H2A/H2B to express an H2B-tdTomato fusion in an upstream direction (H2B promoter) while expressing hph in an downstream direction (H2A promoter). **B**) Representative images from wild type (left), and transformants expressing cytoplasmic hph-tdTomato (plasmid GI3EM18) (center) and nuclear-localized H2B-tdTomato (right). Top row shows DIC and the middle row shows fluorescence microscopy at 561 nm with overlaid images on the bottom row. For comparable results, all strains are presented at the same intensity levels used for H2B-tdTomato fluorescence image. Scale bar indicates ten microns. **C**) Southern blot of four transformants, in which genomic DNA was digested either with XbaI or KpnI and probed using the hygromycin resistance gene (hph). **D**) Four independent transformants were transferred, every two days, in both selective and non-selective medium at 30 °C for a total of 27 days (135-162 mitotic cycles), followed by a challenge on selective media. These strains were spotted in a 2-fold dilution series on non-selective and selective (+Hygromycin) plates, and incubated for 2 days at 30 °C. Image acquisition conditions: POL: transmittance 32%, exposure 0.15 sec; TRITC filter, maximal projection, transmittance 32%, Exposure 0.2 sec, 0.3 micrometers slice thickness.

### Mitotic cycles are fast and highly synchronous during sporogenesis

Chytrid transformation with the divergent construct of H2B-tdTomato resulted in bright nuclear localization of fluorescence when compared to cytoplasmic td-Tomato (Fig 2B). The presence of the T-DNA in the transformants was determined using PCR for hph and H2B-tdTomato (Fig. S4) and confirmed through Southern blot analysis of hph (Fig 2C). The results were consistent with random, single T-DNA genomic integration events in each transformant. Finally, we established that the transformants had transgenerational stability by passaging them in non-selective media for several weeks, followed by a challenge in selective media (Fig 2D).

To quantify the timing and synchrony of the *Spizellomyces* cell cycle, we used live cell epi-fluorescence imaging of H2B-tdTomato strains at 2-minute intervals. Our results show that zoospores have a single nucleus, that the first mitotic event (i.e. one nucleus to two nuclei) occurs after the branching of the germ tube to form a rhizoid, and that sporangia develop and undergo 4-8 mitotic cycles in less than 30 hours before completing their life cycles and releasing 16-256 zoospores; see Movie S1. To measure all nuclei within a sporangium with better temporal and z-resolution, we followed nuclear dynamics at 1-minute time intervals using livecell confocal microscopy of a H2B-tdTomato strain (Movie S2). We measured the number of nuclei over time per sporangia (Fig 3A) to estimate the synchrony of nuclear division waves and the period of time between waves of nuclear division. Wave time (Δt) is the time for a wave of nuclear divisions to propagate across the sporangium. The cell cycle period (*τ*) is the interval of time between nuclear division waves. We found that the average cell cycle period was 150 minutes and that each nuclear division wave is completed within one minute (Fig. 3B). In addition, by following the compaction and localization of H2B-tdTomato we found that all measurable mitotic events occur within 5 minutes, or less than 3.3% of the cell cycle period (Fig. 3B & C)). All together, these results show that *Spizellomyces* is mitotically inactive in its early life cycle (i.e. zoospore, germination). However, once committed, the cell cycle is fast and nuclear divisions are highly synchronous; see Table S1.

**Fig. 3.**
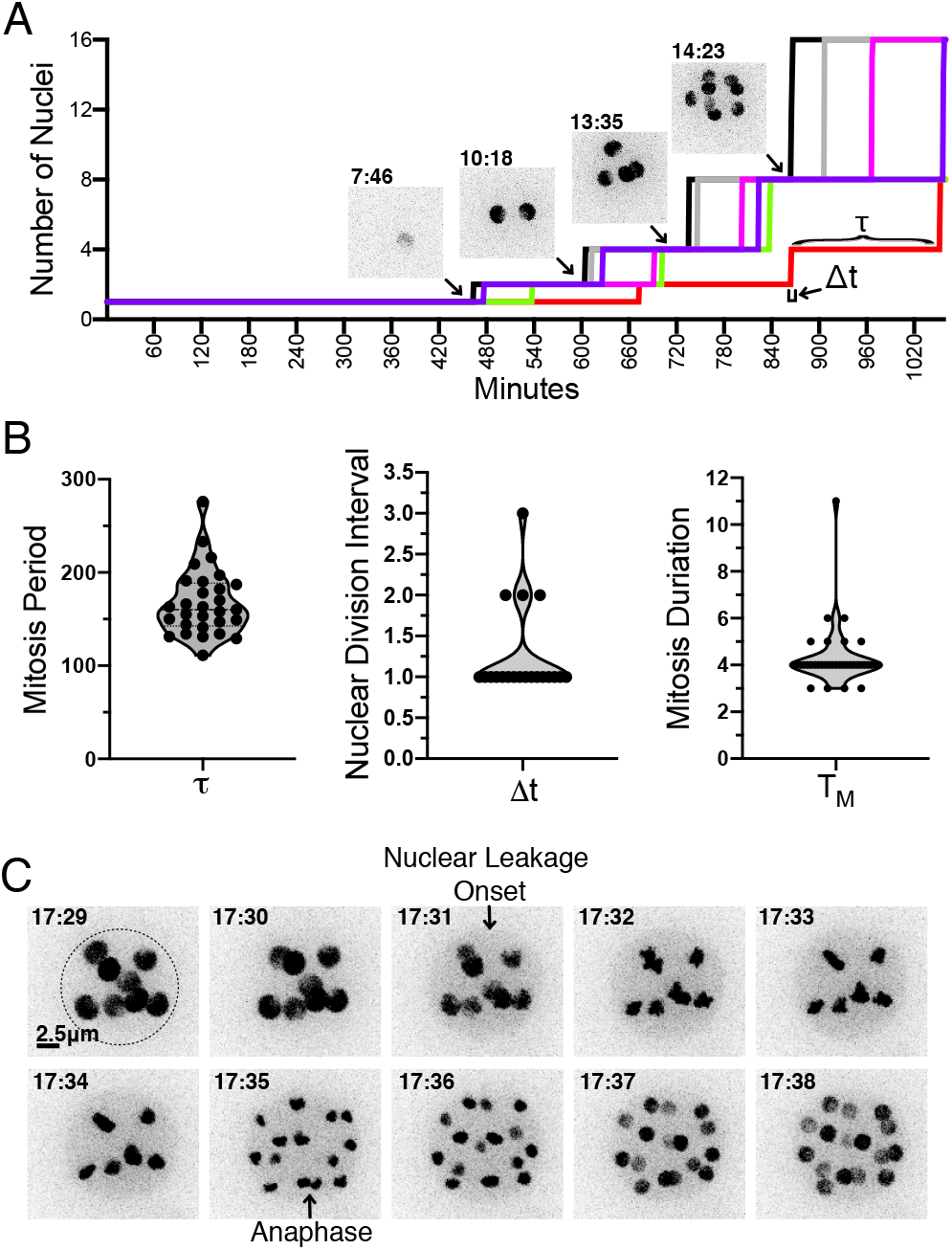
H2B-tdTomato measures the timing and synchrony of mitotic events during sporogenesis. **A)** Number of nuclei as a function of time during the development of a sporangium, along with H2B-tdTomato fluorescence images from select time points. Each colored line corresponds to an independent sporangium. Example of cell cycle period = *τ*, which corresponds to the interval of between doubling of nuclei (i.e. metaphase to anaphase transition) and nuclear division interval (i.e interval of time nuclear number duplicated; Δ*t*). **B)** Distribution of cell cycle period (*τ*; N=29), nuclear division interval (Δ*t*; N=20) and duration of mitosis (*τ*_M_; N=36) across multiple mitotic cycles. **C)** Timing of mitotic events. H2B-tdTomato permits observation of (1) leakage from the nucleus likely due to fenestration of nuclear envelope by the mitotic spindle (8, 21), followed by chromosome condensation, and (2) chromosome separation during anaphase. Dotted line highlights cell wall of sporangium. This particular example shows mitosis duration of 4 min (*τ*_*M*_ = time from nuclear leakage to anaphase). Time in hours:min. Scale 2.5 micrometers.

### Actin polymerization drives zoospore motility

Like their pre-fungal ancestors, chytrids swim with a motile cilium. Some chytrids can also crawl across and between solid substrates, much like amoeba and animal immune cells (8–12, 22–25). Eukaryotes employ multiple strategies to crawl (filopodia, pseudopodia and blebs) that depend on distinct molecular mechanisms (26). One form of crawling, the pseudopod-based α-motility, relies on the expansion of branched-actin filament networks that are assembled by the Arp2/3 complex and allow cells to navigate complex environments at speeds exceeding 20μm/min (12). The activators of branched-actin assembly WASP and SCAR/WAVE have been recently described as a molecular signature of the capacity for α-motility (12). *Spizellomyces* has homologs of WASP (SPPG_00537), SCAR/WAVE (SPPG_02302), and their zoospores are proficient crawlers.

To test whether *Spizellomyces* zoospores crawl using α-motility, we expressed a LifeAct-tdTomato fusion. LifeAct is a 17 amino acid peptide that binds specifically to polymerized actin in a wide variety of cell types, such as actin patches and cables in yeast and actin-filled protrusions of crawling animal cells (27, 28). To confirm that our LifeAct-tdTomato fusion binds specifically to polymerized actin in *Spizellomyces*, we first fixed and stained actin in zoospores (Fig. 4) and sporangia (Fig. 5) with fluorescent phalloidin. We then compared the fluorescent images of cells expressing LifeAct-tdTomato and those expressing hph-tdTomato (negative control) relative to phalloidin-staining.

**Fig. 4.**
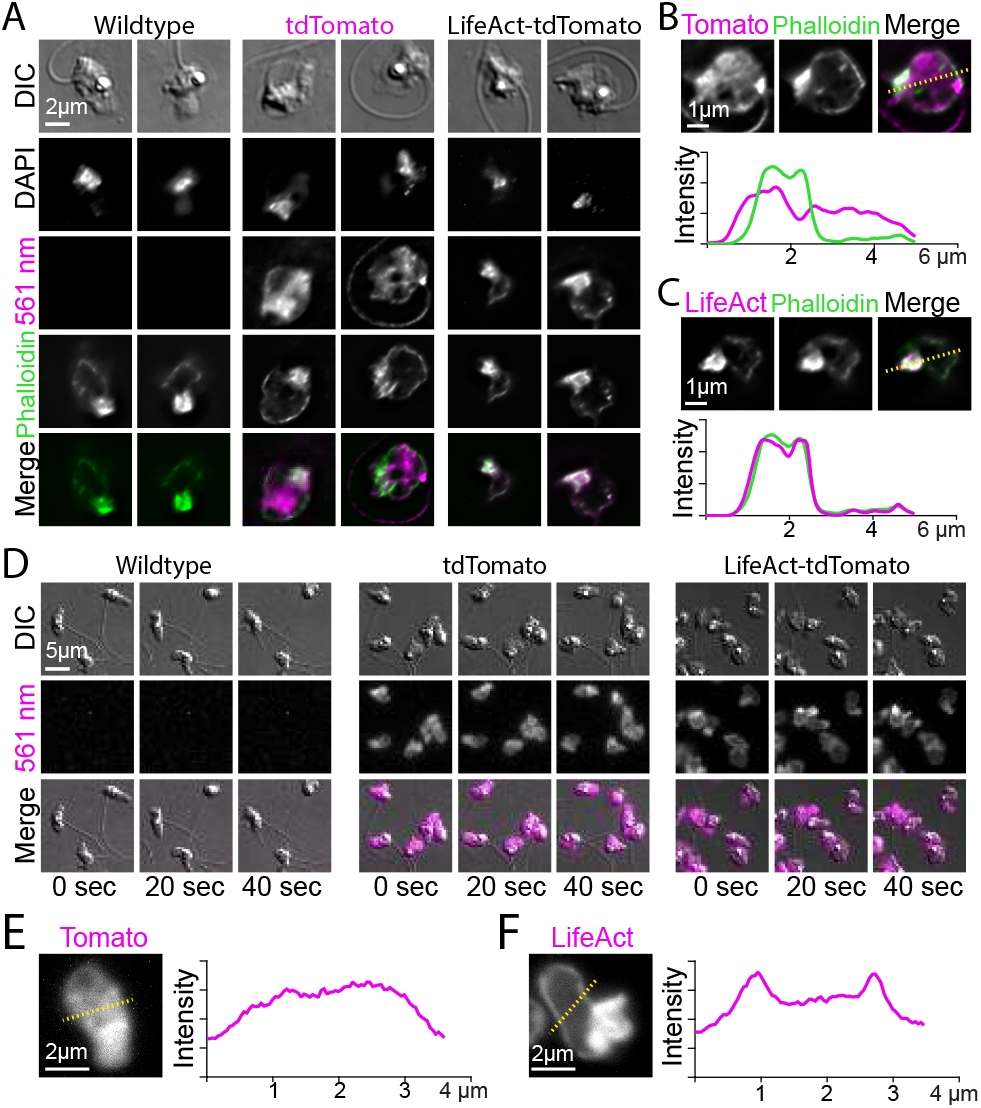
Localization of LifeAct-tdTomato in zoospores highlights cortical and pseudopod actin networks. A) Zoospores from wild type (left), and transformants expressing hph-tdTomato (center) and LifeAct-tdTomato (right) were fixed and stained with fluorescent phalloidin (green). Top row shows DIC and second row shows DNA stain (DAPI). The bottom row shows the phalloidin stain and 561 nm images overlaid. Scale bar indicates two microns. B and C) Line scan of fixed and stained hph-tdTomato (B) and LifeAct-tdTomato fusion (C). The plot shows line scans of normalized fluorescence intensity of the respective fusion protein (magenta) and fluorescent phalloidin (green). The location for generating the line scans is shown by a yellow dotted line in the image above each plot. Scale bars indicate one micron. D) Stills taken at twenty-second intervals from timelapse microscopy of crawling zoospores from the indicated strains at the given timepoints. Images were taken using DIC microscopy (top) and 561 nm fluorescence microscopy (middle), also shown with images merged (bottom). Scale bar indicates five microns. E+F) Line scan of fixed and stained hph-tdTomato (E) and LifeAct-tdTomato fusion (F). The plot shows line scans of normalized fluorescence intensity of the respective fusion protein (magenta) and fluorescent phalloidin (green). The location for generating the line scans is shown by a yellow dotted line in the image above each plot. Scale bars indicate two microns. See also Movie S3.

**Fig. 5.**
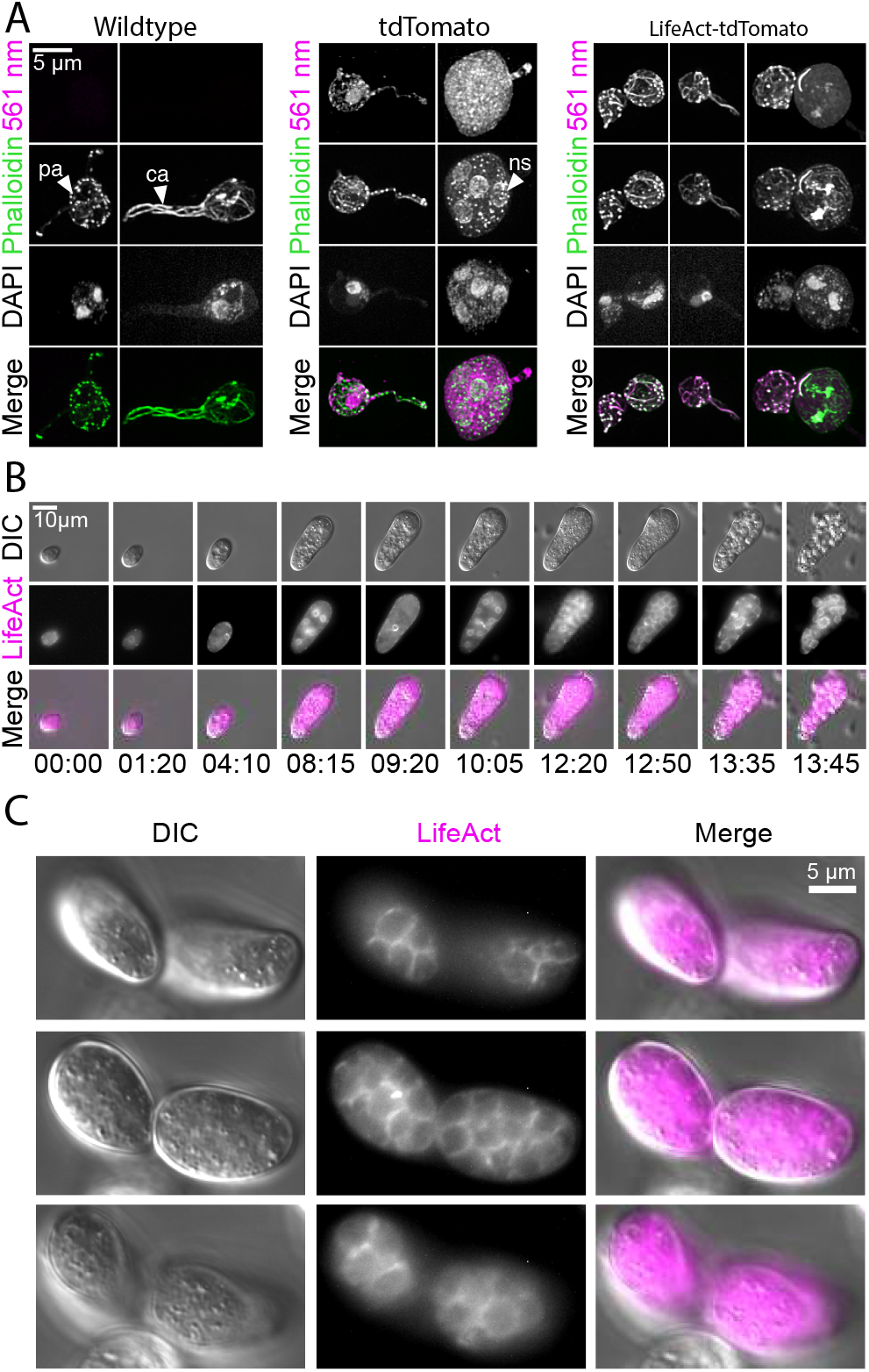
Localization of LifeAct-tdTomato in sporangia highlights actin patches, cables, and perinuclear shells. A) Sporangia from wild type (left), and transformants expressing hph-tdTomato (center) and LifeAct-tdTomato (right) were fixed and stained with fluorescent phalloidin (green). The third row shows DNA stain (DAPI). The bottom row shows the phalloidin stain and 561 nm images overlaid. Scale bar indicates five microns. Arrows point to examples of actin patches (“pa”), cables (“ca”) and perinuclear actin shells (“ns”). B) Selected stills taken from time-lapse microscopy of developing sporangia from the LifeAct-tdTomato transformant strain at times indicated (hours: minutes). Formation of polygonal territories (“pt”) precedes cellularization. Images taken using DIC (top) and 561 nm (middle), also shown with images merged (bottom). Scale bar indicates ten microns. See also Movie S4. C) Multiple planes of a single sporangium show how polygonal territories formed during later stages of cellularization encompass all cytoplasm.

In contrast to the relatively homogeneous distribution of fluorescence in hph-tdTomato zoospores, both fixed (Fig. 4A-C) and living zoospores (Fig. 4D-E, and Movie S3) of the LifeAct-tdTomato strain showed a thin layer of fluorescence at the cell cortex, and high levels of fluorescence in the pseudopods at the leading edge. Because the LifeAct fusion localization nearly perfectly correlated with phalloidin intensities in fixed cells, but the hph fusion did not, we presume that this fluorescence represents polymerized actin. This distribution of actin is in agreement with α-motility and actin localization observed for fixed cells during zoospore crawling of *Neocallimastix* (29) and *Batrachochytrium* (12).

### Actin polymerization during sporogenesis

Once the chytrid zoospore encysts, it builds a sporangium by growing both radially and in polarized fashion during germ tube extension and rhizoid formation (Fig. 1). During early sporo-genesis, the nuclei are very dynamic while replicating and dividing but then stop moving in late sporogenesis, presumably during cellularization and zoospore formation (Movie S1). Actin has been reported to play fundamental roles during cellularization in another chytrid *Allomyces macrogynus* (30). Thus, we expect polymerized actin to play a role in the nuclear dynamics and cellularization during sporogenesis in *Spizellomyces*.

Our experiments revealed an actin cytoskeleton distributed primarily between cortical patches and linear structures that resemble the actin cables of other fungal species (Fig. 5A). Each of these structures was visible in both fixed phalloidin stained cells and living cells expressing the LifeAct-tdTomato gene. We also detected transient perinuclear polymerized actin shells with a period similar to mitotic events (Fig. 5B and Movie S4), which suggests a role for polymerized actin during nuclear division. In late sporogenesis, polymerized actin delineated polygonal zoospore territories (Fig. 5B and 5C). This is reminiscent of epithelial cellularization seen during *Drosophila* embryogenesis when syncytial nuclei are encapsulated by cell membrane.

## Discussion

Here, we report stable and robust genetic transformation of the chytrid *Spizellomyces punctatus*. We identified and tested native *Spizellomyces* promoters, including a divergent H2A/H2B promoter that can simultaneously express hygromycin resistance (hph) and a gene of interest throughout the chytrid life cycle. This design has been successfully used to express fluorescent proteins and resistance genes in Dikaryotic marine fungi (data not shown), which suggests they should be useful for transforming other chytrids, such as the pathogenic *Batrachochytrium*. Agrobacterium was previously used to transform the aquatic chytrid *Blastocladiella emersonii* (20). Unfortunately, genetic transformation of *Blastocladiella* displayed a narrow window of gene expression and could only be detected at the zoosporic stage. The advances described in this study should be useful for random and possibly targeted gene integration in other chytrids.

As a first step to characterize the chytrid cell cycle, we used an H2B-tdTomato fusion and live-cell microscopy to measure the timing of nuclear division. The *Spizellomyces* developmental program allocates a narrow window (5min) of time to the mitotic process during each cell cycle with a highly synchronous wave of nuclear divisions. The level of synchrony is similar to plasmodial nuclei in *Physarum polycephalum* or syncytial nuclei during *Drosophila* development, where all nuclei divide within 2 min (31) (see Table S1). These mitotic dynamics make *Spizellomyces* an interesting comparative model for exploring the physical and molecular determinants of cell division synchrony and its evolution in fungi, animals, and amoeba.

By visualizing actin dynamics throughout the chytrid life cycle we found that *Spizellomyces* zoospores assemble thick actin cortexes and build actin-filled protrusions during motility, similar to other chytrid species (12), animal cells, and amoebae. Once the zoospores encyst, there is a drastic shift in actin cytoskeleton organization, in which the cortical shell of polymerized actin was replaced by dynamic puncta (actin patches), some of which are associated with actin cables that often extended into the germ tube or rhizoids. This architecture is typical of fungi, where actin patches are associated with endocytosis and cell wall deposition, where as actin cables are pathways for targeted delivery of exocytic vesicles (32). This biphasic actin distribution – an actin cortex and actin-filled protrusions in zoospores, and actin patches and cables in sporangia – indicates that, like its cell cycle regulatory network (13), the actin cytoskeleton of *Spizellomyces* displays features that resemble those of both animal and fungal cells.

Our results revealed further actin dynamics and organization during the chytrid development. This includes the formation of perinuclear actin shells that are fleetingly detected by live imaging. Previous observations suggested that perinuclear actin shells of anaerobic chytrids were a fixation artifact (29); however, our live cell data indicate that these are real, dynamic cellular structures that likely occur in many chytrids. Similar perinuclear actin shells are associated with nuclear lamina in animal cells, where they play a role in changing nuclear shape before and after mitosis (33, 34) or when squeezing through narrow channels (35). Although chytrids and other fungi have lost nuclear lamins, it seems likely that *Spizellomyces* perinuclear actin shells are associated with changes in nuclear shape during the cell cycle.

Finally, we showed that polymerized actin is likely involved in the formation of zoospore polygonal territories (Fig. 5b) before the formation of the cleavage planes during cellularization. In contrast to cellularization in *Drosophila*, which occurs at the surface of the embryo (i.e. twodimensional cellularization), cellularization in *Spizellomyces* happens in a three-dimensional context within three hours. The chytrid cellularization process is reminiscent of the polarized epithelium of the social amoeba *Dictyostelium discoideum* (36, 37) and the membrane invagination dynamics during cellularization in the ichthyosporean *Sphaeroforma arctica*, an unicellular relative of animals (38). To what extent is the emergence of multicellularity, which appears to have evolved independently in amoeba, animals, and fungi, dependent upon a shared ancestral toolkit? Establishing shared and unique traits between chytrid fungi and other key lineages will provide a powerful cross-lineage experimental system to test core hypotheses on the evolution of multicellularity and derived fungal features. Based on ouyr tools and finding, we expect that *Spizellomyces* will be a useful model system to study the evolution of key animal and fungal traits.

## Acknowledgments

This material is based upon work supported by the National Science Foundation under Grants No. IOS-1827257 (L.K.F.-L.) and IOS-1755248 (N.E.B.). Spinning disk confocal microscopy data collection of fixed cells was performed in the Light Microscopy Facility and Nikon Center of Excellence at the Institute for Applied Life Sciences, University of Massachusetts Amherst with support from the Massachusetts Life Science Center. The authors are grateful to Joe Heitman for his unwavering support throughout the project and to David Booth, Stefano Di Talia, and Danny Lew for comments on the manuscript.

## Materials & Methods

### Strains and growth conditions

We used *Spizellomyces punctatus* (Koch type isolate NG-3) Barr (ATCC 48900) for all chytrid experiments. Unless otherwise stated, *Spizellomyces* were grown at 30°C in Koch’s K1 medium (1L; 0.6g peptone, 0.4g yeast extract, 1.2g glucose, 15g agar if plates) (39). Two days prior to harvesting zoospores, we aliquoted and spread 1mL of active, liquid culture pregrown in K1 medium onto K1 plates and incubated them to allow zoospores to encyst, mature, and colonize the agar surface. We flooded each active *Spizellomyces* plate with 1 mL of dilute salt (DS) solution (40) and incubated at room temperature. After one hour, released zoospores were retrieved by harvesting the DS medium and purified by slowly filtering the harvest in Luer-Lok syringe through an autoclaved syringe filter holder (Advantec REF:43303010) preloaded with a 25mm Whatman Grade 1 filter paper (CAT No. 1001-325).

### Plasmids

We used the pGI3 plasmid backbone for Agrobacterium-mediated transformation (19), which contains the *Saccharomyces cerevisiae* 2μ replication origin and the URA3 selectable marker. This allows pGI3 and its derivatives to replicate in *E. coli*, *A. tumefasciens* and *S. cerevisiae*. Complete details of primers and plasmid construction are in Supplementary Materials.

### Agrobacterium-mediated transformation of *Spizellomyces*

We prepared competent *Agrobacterium* EHA105 strains following the protocol of (41). Plasmids were transformed into competent *Agrobacterium* using 0.2 cm cuvettes in a Gene Pulser electroporator (Bio-Rad, USA) at 25 μF, 200 Ω, 2.5kV. Single colonies were streaked on selective plates (Kanamycin). A colony of transformed *Agrobacterium* containing pGI3-derived plasmid was grown overnight at 30°C in 5 mL of Luria-Bertani broth supplemented with Kanamycin (50mg/L). After centrifugation, the cell pellet was resuspended in 5mL of IM (16), diluted to an OD_660_ of 0.1 and grown under agitation at 30°C until achieving a final OD_660_ of 0.6, at which point the culture was ready for co-culturing with chytrid (300μL per transformation). IM; composed of MM salts (42) and 40M 2-(N-morpholino)ethanesulfonic acid (MES), pH 5.3, 10 mM glucose, 0.5% (w/v) glycerol and 200μM Acetosyringone);

In parallel, chytrid zoospores were harvested and pelleted by centrifugation at 800g for 10 min. Zoospores were then gently resuspended in 300μL of induction medium (IM). We found that one K1 plate provides enough zoospores for one transformation. For every transformation, zoospores and *Agrobacterium* were combined at four different ratios; 1:1, 1:0.25, 0.25:1, and 0.25:0.25 in a total volume of 200μL. The surface of the IM plate was rubbed with the bottom of a sterile glass culture tube to generate slightly concave depressions in each quadrant of the plate. Each 200μL co-incubation mixture was then plated and incubated unsealed for 4 days at room temperature. Mock transformations with empty *Agrobacterium* (no binary plasmid; grown in the absence of plasmid selective medium) were included as a negative control.

After co-incubation, we added 1mL of DS solution and gently scraped the plate with a razor blade, pooling the different cell ratios into a single 50mL centrifuge tube, raising the volume to 20mL with DS solution, and re-suspending clumps by inversion. The mixture was centrifuged at 1000g for 10 min and the liquid phase was discarded. The remaining pellet was carefully resuspended with DS solution, plated on K1 plates containing Ampicillin (50mg/L) and Tetracycline (50mg/L) to select against *Agrobacterium* and Hygromycin (200mg/L) to select for transformed *Spizellomyces*. *Spizellomyces* survival controls were performed by plating transformations after co-culture in non-selective K1 media (Ampicillin (50mg/L), Tetracycline (50mg/L)). Transformation plates and controls were incubated at 30C until colonies were observed (5-6 days). All plates were sealed with parafilm to prevent desiccation. Single colony isolates were retrieved with a sterile needle, resuspended in DS solution and re-plated on a selective Hygromycin plate.

### Nucleic acid manipulation

High molecular weight genomic DNA extraction was performed using CTAB/Chloroform protocol (CTAB/PVP buffer: 100mM Tris-HCl pH7.5; 1.4M NaCl; 10mM EDTA; 1% CTAB; 1% PVP; 1% β-Mercaptoethanol (added just before use)). Briefly, 10 plates of the selected strain were grown at 30°C for two days and zoospores were harvested and purified as described before. The pellet of zoospores was resuspended directly in 900μL of CTAB/PVP buffer pre-warmed to 65°C and transferred to 2mL centrifuge tubes. Tubes were centrifuged briefly and incubated at room temperature for one hour in a nutating mixer. After 5min incubation on ice, DNA was extracted with Chloroform (Sigma-Aldrich REF:288306) twice followed by treatment of supernatant with 100ng RNase A (Biobasic; 60U/mg Ref:XRB0473) for 30min at room temperature. DNA was then precipitated by adding 0.2 volume of 10M Ammonium acetate and one volume of absolute Isopropanol and incubated at 4°C overnight. DNA pellet was washed with 70% ethanol thrice and resuspended in TE buffer. DNA quality and concentration was determined by gel electrophoresis.

Detection of transgene integration by Southern blot. 1μg of high-molecular weight gDNA from each strain was treated overnight at 37°C with 10U of XbaI or KpnI-HF restriction enzymes, resolved on a 1% Agarose 1X TAE (Tris-Acetate-EDTA) gel and blotted to a GE Healthcare Amersham Hybond-N+ membrane. The membrane was hybridized with a 809bp fragment of the Hygromycin resistance gene amplified with primers HygF1 and HygR1 and radiolabeled with [α-^32^P]-dCTP using the Prime-It II Random Primer Labelling Kit (Agilent Technologies; REF:300385) following manufacturer’s instructions.

### Microscopy

H2B-tdTomato-expressing chytrids were harvested from plates, placed in a glass-bottom dish (Mattek), and covered with a 1.5% K1 agarose pad to keep cells healthy and in the plane of focus (43). Our timelapse movies harvested zoospores from plates and re-suspended them in Leu/Lys paralyzing solution (44) before putting them in glass-bottom dishes, as above. Live-cell epifluorescence was performed on a temperature-controlled Deltavision Elite inverted microscope equipped with 60x/1.42 oil objective and Evolve-512 EMCCD camera. Excitation light was 542/27 nm (7 Color InsightSSI) and emission to the camera was filtered by 594/45 nm (TRITC). Epifluorescence live-cell imaging was done at 30 C. Live-cell confocal microscopy was performed on a Nikon Eclipse Ti inverted microscope equipped with a 100x/N.A. 1.49 CFI Apo TIRF oil objective and fitted with a Yokogawa CSU-X1 spinning disk and Andor iXon 897 EMCCD camera. Excitation light was via 488 nm laser. Confocal live-cell imaging was done at 27 C.

LifeAct-expressing zoospores were collected in DS solution and transferred to cover-glass bottom dishes. For phalloidin staining, glass coverslips were plasma cleaned and immediately coated with 0.1% polyethyleneimine for 5 minutes, washed thrice with water, then overlaid with zoospores or sporangia suspended in DS solution. Cells were allowed to adhere for 5 minutes before fixation by adding four volumes of 4% paraformaldehyde in 50mM Cacodylate buffer (pH 7.2). Cells were fixed for 20 minutes on ice, washed once with PEM buffer (100 mM PIPES, 1 mM EGTA, 0.1 mM MgSO_4_), permeabilized and stained with 0.1% Triton X in PEM with 1:1000 Alexa Fluor 488 Phalloidin (66 nM in DMSO, Sigma D2660) for 10 minutes at room temperature, washed with PEM, then mounted onto glass slides using Prolong Gold with DAPI (Invitrogen P36931). Zoospores were imaged on a Nikon Ti2-E inverted microscope equipped with 100x oil PlanApo objective and sCMOS 4mp camera (PCO Panda). Excitation light was via epi fluorescence illuminator at 405 nm, 488 nm, and 561 nm. Sporangia were imaged on a Nikon Ti-E inverted microscope equipped with 100X oil objective and fitted with a Yokogawa X1 spinning disk (CSU-W1) with 50 μm pinholes and Andor xIon EMCCD camera. Excitation light was via 405 nm laser, 488 nm laser, and 561 nm laser. Z-stack fluorescent images were deconvolved with NIS Elements v5.11 using 20 iterations of the Richardson-Lucy algorithm. Image analysis was performed with the ImageJ bundle Fiji (45). All imaging was done at room temperature.

## Supplementary Material

### Plasmid construction

We initially built base plasmids and later cloned parts into the T-DNA of the Agrobacterium pGI3 plasmid. pGI3 is a binary plasmid derived from pPZP201-BK (KanR) (18) and pRS426, which contains the *Saccharomyces cerevisiae* 2μ origin of replication and the URA3 selectable marker. This let us screen gene expression and GFP fluorescence from pGI3EM plasmids transformed into *Saccharomyces*. We identified *H2A* (SPPG_02344), *H2B* (SPPG_02345), and *HSP70* (SPPG_04820) genes and promoters from FungiDB (46) by blasting human homologs against the *Spun* genome. The promoter DNA was cloned from *Spizellomyces* genomic DNA (gDNA) isolated using the methods described in Materials & Methods. Coding DNA was cloned from a cDNA library of *Spizellomyces* transcripts. RNA was extracted from a mixed population *Spizellomyces* zoospores and sporangia using Qiagen RNeasy Plant Mini Kit following manufacturer procedures and RLT buffer. cDNA was synthetized using Thermo Scientific Maxima H Minus Strand cDNA synthesis Kit with dsDNase (REF:K1682) following manufacturer instructions with oligo (dT)_18_ primers.

#### Base plasmids

- pEM01 (CMVpr-hph) was constructed by digesting pNB780 (a pRS406 plasmid) with SacI-HF and BglII, isolating the large backbone fragment, and then assembling CMVpr (primers CMV_F and CMV_R), hph (primers Hyg_F and Hyg_R) and ADH1 terminator (primers ADH1t_F and ADH1t_R) in a single Gibson reaction following manufacturer instructions (NEB, E2611S). CMVpr was obtained by PCR from pNB419 (pAB1T7 = CMVpr-TetR-GFP-VP16), hygromycin resistance gene (hph) from pRS306H, and *Saccharomyces cerevisiae* ADH1 terminator from pNB780. The final plasmid was verified by analytical restriction digest and Sanger sequencing (primers pRS-up and ADH1t-dn).
- pEM03 (Hsp70pr-hph) was constructed by digesting pEM01 with SacI and PacI, isolating the large backbone fragment, and then inserting Hsp70pr using Gibson cloning. Hsp70pr was obtained by PCR from *Spizellomyces* gDNA using primers HSP70_F and HSP70_R. The final plasmid was verified by analytical restriction digest and Sanger sequencing (primers pRS-up and ADH1t-dn).
- pEM09 (Hsp70pr-hph-GFP) was constructed by digesting pEM03 with PacI and AscI, isolating the large backbone fragment, and then inserting hph-GFP using Gibson cloning. Hygromycin resistance gene (hph) was obtained by PCR from pFA6-GFP(S65T)::hph using primers HygR_F and HygR_R, whereas GFP(S65T) was amplified from the same plasmid using primers GFP_F and GFP_R. The final plasmid was verified by analytical restriction digest and Sanger sequencing (primers pRS-up and ADH1t-dn).

#### Agrobacterium plasmids

- pGI3EM09 (Hsp70pr-hph-GFP) was constructed by digesting pGI3 with HindIII and EcoI, isolating the large backbone fragment, and then inserting Hsp70pr-hph-GFP in the multiple cloning site between the T-DNA LB and RB borders of pGI3 using SLIC cloning (47, 48). Hsp70pr-hph-GFP was obtained by PCR of pEM09 using primers GI3EM9IIF and GI3EM9IIR. The final plasmid was verified by analytical restriction digest and Sanger sequencing (Standard primers M13F and M13R).
- pGI3EM11 (H2Bpr-hph-GFP) was constructed by digesting pGI3EM09 with HindIII and PacI, isolating the large backbone fragment, and then inserting H2Bpr using SLIC cloning. H2Bpr was PCR amplified from *Spizellomyces* gDNA using primers prH2B_F and prH2B_R. The final plasmid was verified by analytical restriction digest and Sanger sequencing (Standard primers M13F and M13R).
- pGI3EM18 (H2Bpr-hph-tdTomato) was constructed by digesting pGI3EM11 with HindIII and AscI, isolating the large backbone fragment, and then re-inserting PCR amplicon of H2Bpr-hph with an extra BamHI site 3’ (primers prH2B_F and GI3EM11up) and tdTomato using SLIC cloning. TdTomato was obtained by PCR of pKT356 (pFA6a-tdTomato::SpHIS5; gift from Daniel J. Lew) using primers HygtdTom_F and AdhtdTom_R. The final plasmid was verified by analytical restriction digest and Sanger sequencing (Standard primers M13F and M13R).
- pGI3EM29 (H2Bpr-hph-mCitrine) was constructed by digesting pGI3EM18 with BamHI and AscI, isolating the large backbone fragment, and then inserting mCitrine using SLIC cloning. mCitrine was obtained by PCR of mCitrine-PCNA-19-SV40NLS-4 (Addgene plasmid # 56564), a plasmid created by the Davidson lab (49). The final plasmid was verified by analytical restriction digest and Sanger sequencing (Primers M13 and H2BprF2).
- pGI3EM30 (H2Bpr-hph-mClover3) was constructed by digesting pGI3EM18 with BamHI and AscI, isolating the large backbone fragment, and then inserting mClover3 using SLIC cloning. mClover3 was obtained by PCR of pKK-mClover3-TEV (Addgene plasmid # 105778), a plasmid created by the Dziembowski lab (50). The final plasmid was verified by analytical restriction digest and Sanger sequencing (Primers M13 and H2BprF2).
- pGI3EM31 (H2Bpr-hph-mCerulean3) was constructed by digesting pGI3EM18 with BamHI and AscI, isolating the large backbone fragment, and then inserting mCerulean3 using SLIC cloning. mCitrine was obtained by PCR of mCerulean3-N1 (Addgene plasmid # 54730), a plasmid created by the Davidson lab (49). The final plasmid was verified by analytical restriction digest and Sanger sequencing (Standard primers M13F and M13R).
- pGI3EM20B (H2Apr-hph::H2Bpr-H2B-TdTomato) took advantage of a divergent *Spizellomyces* H2A (SPPG_02344) and H2B (SPPG_02345) promoter to express an H2B-TdTomato fusion in one direction (H2B promoter) and hph in the other direction (H2A promoter). This plasmid contained the H2B gene with introns. pGI3EM20B was constructed by digesting pGI3EM18 with HindIII and BamHI, isolating the large backbone fragment, and then assembling hph (from pEM03, primers AdhHygF and AdhHygR), H2A/H2Bpr (primers H2B2D_F and H2B2D_R) and the H2B gene (primers H2Bgen_F and H2Bgen_R) using a four piece SLIC assembly. The final plasmid was verified by analytical restriction digest and Sanger sequencing (Standard primers M13F, M13R and primer H2BprF1).
- pGI3EM20C (H2Apr-hph::H2Bpr-H2B-TdTomato) is a version of pGI3EM20B, where the intron of H2B has been removed. pGI3EM20C was constructed by digesting pGI3EM20B with SalI and BamHI, isolating the large backbone fragment, amplifying the coding version of H2B from *Spizellomyces* cDNA using the same primers (primers H2Bgen_F and H2Bgen_R) followed by SLIC cloning. The final plasmid was verified by analytical restriction digest and Sanger sequencing (Standard primers M13F, M13R, and primer H2BprF1).
- pGI3EM22C (H2Apr-hph::H2Bpr-Lifeact-TdTomato) was constructed by digesting pGI3EM20C with SalI and BamHI, isolating the large backbone fragment, and then inserting LifeAct using standard ligation cloning. LifeAct is a 17 amino acid peptide that binds specifically to polymerized actin in a wide variety of cell types, such as actin patches and cables in yeast and actin-filled protrusions of crawling animal cells (27, 28). LifeAct tag was synthesized as a 300 bp gene fragment through Twist Bioscience in which LifeAct-G(4)S was flanked upstream by SalI restriction site and H2Bpr 5’UTR and downstream by BamHI restriction site and part of the 5’ tdTomato sequence. This gene fragment was amplified by PCR, digested by SalI and BamHI and then ligated into pGI3EM20C backbone fragment. The final plasmid was verified by analytical restriction digest and Sanger sequencing (primer H2BprF2).

**Movie S1**: Time-lapse microscopy of nuclear divisions in developing H2B-tdTomato sporangia taken over the course of 24 hours with images captured every 2 minutes. The left panel shows 561 nm fluorescence microscopy of H2B-tdTomato and the right panel shows DIC microscopy. Time-stamp is hr:min. Scale bar indicates ten microns.

**Movie S2**: Time-lapse microscopy of nuclear divisions in developing H2B-tdTomato sporangia taken over the course of 24 hours with images captured every 1 minute. Time-stamp is hr:min. Scale bar indicates five microns.

**Movie S3**: Time-lapse microscopy of crawling LifeAct-tdTomato zoospores taken over the course of two minutes with images captured every second, playback in real time. The left panel shows DIC microscopy, the center panel shows 561 nm fluorescence microscopy, and the right panel shows a merge of fluorescence and DIC microscopy. Time stamp is min:secs. Scale bar indicates ten microns.

**Movie S4**: Time-lapse microscopy of developing LifeAct-tdTomato sporangia taken over the course of 19 hours with images captured every 5 minutes. The right panel shows DIC microscopy, the center panel shows 561 nm fluorescence microscopy, and the right shows a merge of fluorescence and DIC microscopy. Time stamp is hr:mins. Scale bar indicates ten microns.

**Table S1.** Comparison of nuclear division synchrony for different coenocytic organisms. Wave time (Δt) is defined as the average interval of time for a wave of synchronized nuclear divisions to propagate across the coenocytic nuclei. The nuclear division period (*τ*) is the average interval of time between waves. Organisms are listed from highest to lowest synchrony index.

| Coenocytic organism | Length scale<br>($\mu m$) | Wave Time<br>$\Delta t$ (min) | Wave Speed<br>($\mu m \text{ min}^{-1}$) | Nuclear Division<br>Period $\tau$ (min) | Synchrony index<br>$100\% \cdot (1 - \Delta t / \tau)$ | References |
| --- | --- | --- | --- | --- | --- | --- |
| *Physarum polycephalum* (amoeba) | 1000 | 2 | 500 | 840 | 99.8% | (51) |
| *Spizellomyces punctatus* (fungi) | 5-10 | 1 | 5-10 | 150 | 99.3% | This work |
| *Creolimax fragrantissima* (holozoa) |  | 20 |  | 300 | 93.3% | (52) |
| *Drosophila melanogaster* (metazoa) | 500 | 1.5 | 360 | 10 | 85.5% | (31) |
| *Aspergillus nidulans* (fungi) | 700 | 20 | 35 | 60 | 66.7% | (53, 54) |

**Table S2.**
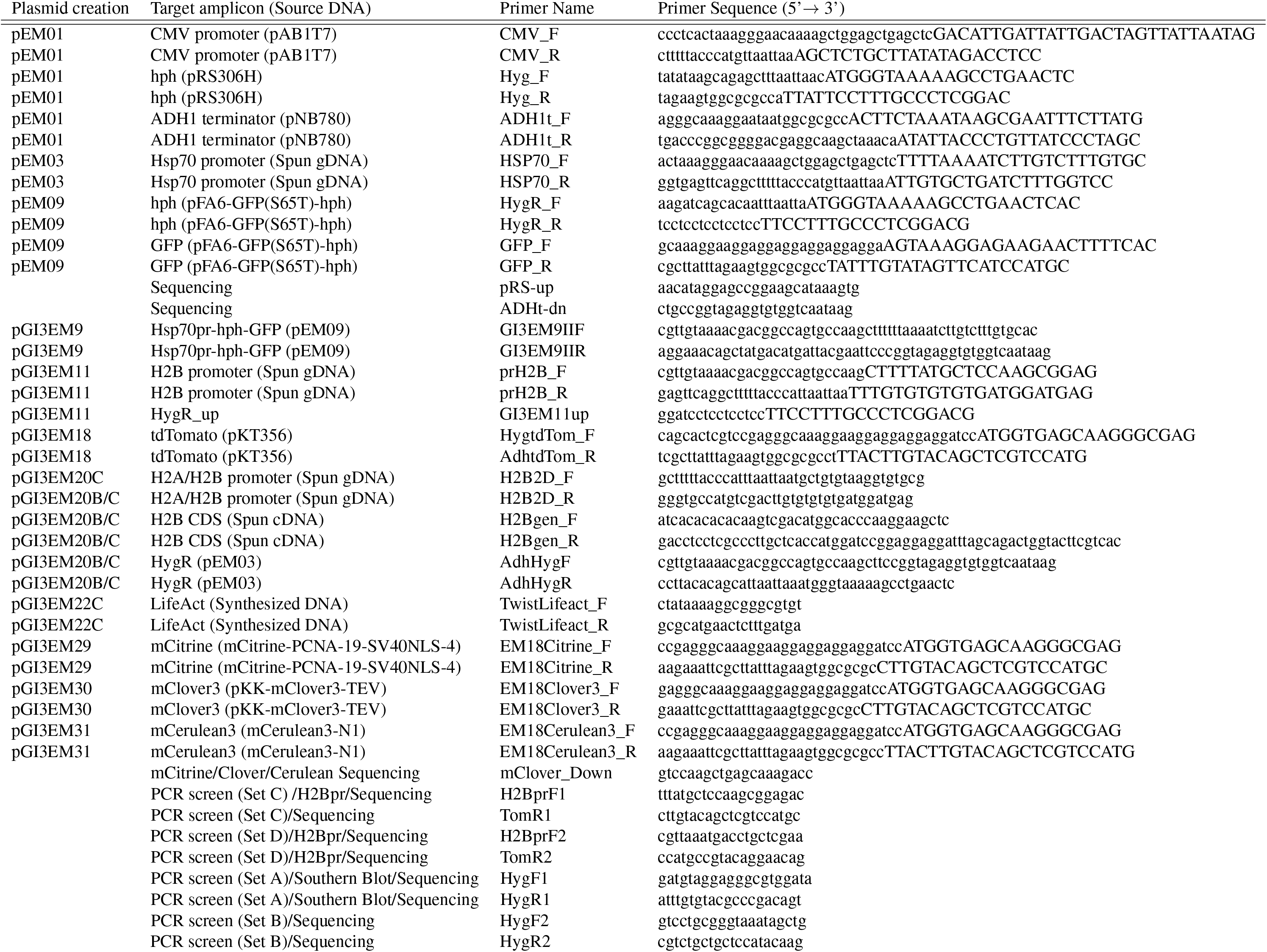
Primers used in this study. Capital letters in SLIC and Gibson primers indicate template binding regions.

**Fig. S1.**
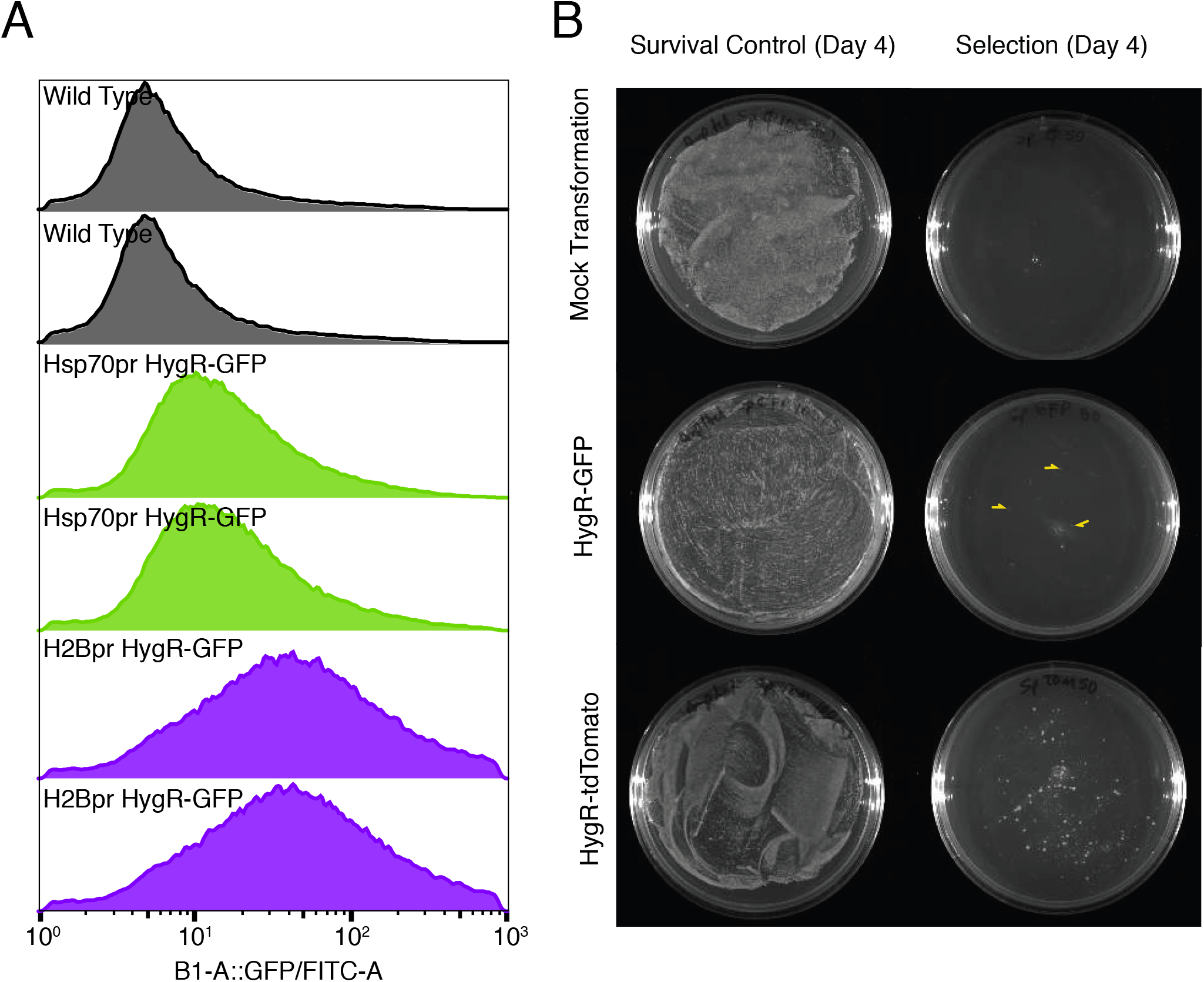
Spizellomyces promoters successfully express fluorescent proteins and drug-resistance genes. **A)** Fluorescence distribution of yeast strains transformed with plasmids pGI3EM09 (Hsp70pr-hph-GFP) and pGI3EM11 (H2Bpr-hph-GFP) relative to untransformed wild-type (WT). Flow cytometry data were collected on a MacsQuant VYB with a 488 nm excitation laser and FITC emission filter (525/50 nm). Data were collected from two independent transformants. The large width of the fluorescence distribution arises from copy number fluctuations of the 2μ plasmid in *Saccharomyces cerevisiae*. **B)** Typical time scale and transformation efficiency using Agrobacterium-mediated transformation of *Spizellomyces*. Data is shown for pGI3EM11 (H2Bpr-hph-GFP) and pGI3EM18 (H2Bpr-hph-tdTomato) in the absence and presence of selection (200 mg/L Hygromycin). Ampicillin and tetracycline (50 mg/L) are included to kill any *Agrobacterium* transferred from the co-culture plate; see Materials & Methods. Small yellow triangles indicate three examples of tiny colonies that appeared on pGI3EM11 plate on Day 4.

**Fig. S2.**
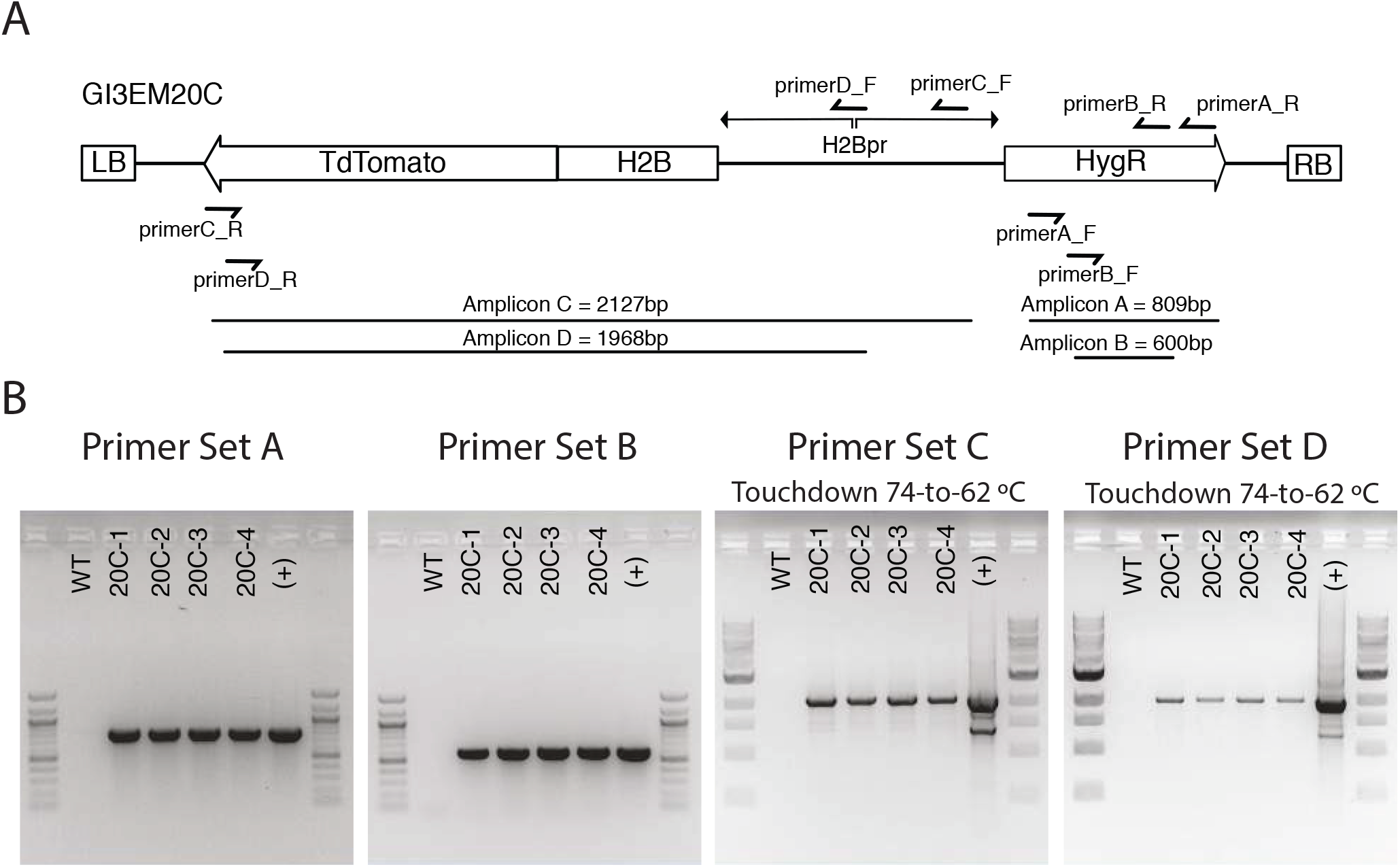
PCR validation of H2B-tdTomato transformants. **A)** Primer locations and amplicon sizes of different pGI3EM20C primer pairs. **B)** Gel electrophoresis of different PCR reactions using genomic DNA of untransformed (WT) and four independent *Spizellomyces* transformants. (+) control used pGI3EM20C plasmid DNA as template.

**Fig. S3.**
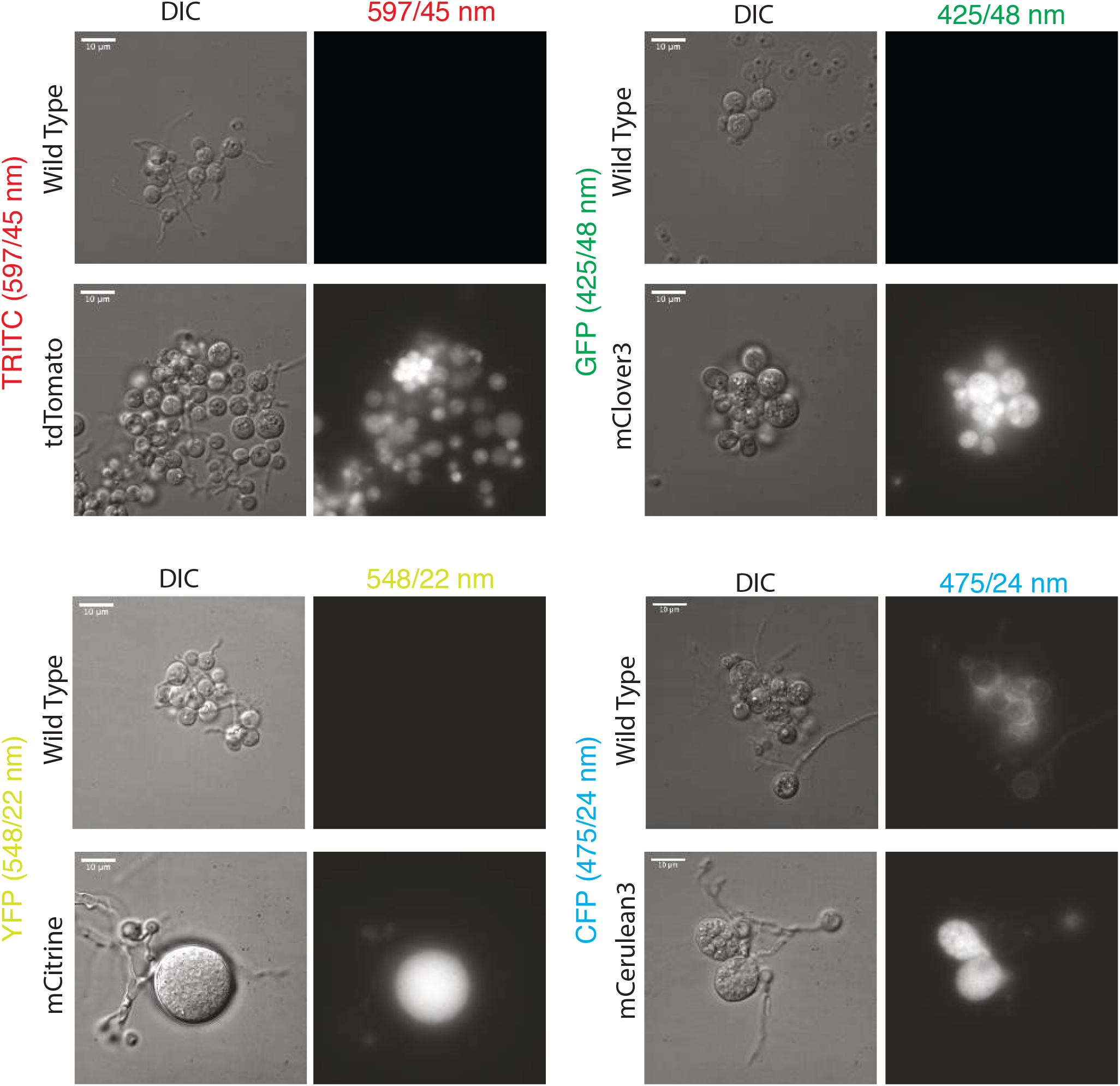
Diverse fluorescent proteins are functional in *Spizellomyces punctatus*. DIC and fluorescence images were taken with a Deltavision Elite microscope using POL Transmission 32%, exposure time 0.5sec; Filter Transmission 32%, exposure time 0.3sec. TRITC filter set excitation (542/27 nm) and emission (597/45 nm); GFP filter set excitation (475/28 nm) and emission (525/48 nm); YFP filter set excitation (513/17 nm) and emission (548/22 nm); CFP filter set excitation (438/24 nm) and emission (475/24 nm). Paired wild-type and fluorescent protein strains are plotted with the same intensity range.

